# Pure CD32+ CD4+ Cells Are Cytotoxic Memory CD4+ T Lymphocytes Highly Enriched for HIV-1 DNA

**DOI:** 10.1101/2025.07.09.663884

**Authors:** Philipp Adams, Elena Herrera-Carrillo, Alexandra Vujkovic, Aldo Jongejaan, Perry M. Moerland, Majdouline El Moussaoui, Gilles Darcis, Ben Berkhout, Alexander O. Pasternak

**Author notes:** Corresponding author. Mailing address: AMC, Room K3-113B, Meibergdreef 15, 1105 AZ Amsterdam, The Netherlands.

## Abstract

The elusive viral reservoir is the major obstacle to an HIV-1 cure. Cell surface protein CD32 has been proposed to pinpoint cells with a very high proviral enrichment, potentially providing a tool for selective therapeutic targeting of the reservoir. However, this finding has been challenged by subsequent reports. In order to clarify the nature of CD32 expression on CD4+ T cells, here we designed a purification strategy allowing single-cell analysis using state-of-the-art technologies. The established multilevel sorting strategy allowed to characterize a rare (median 0.06% of CD4+ T cells) but bona fide CD32+ CD4+ T-cell population. In vitro cell stimulation could not lead to CD32 upregulation. Detailed phenotyping revealed that CD32+ T cells reside mostly in the memory T-cell compartment with heightened levels of acute and chronic activation markers. Single-cell RNA sequencing identified that CD32+ CD4+ T cells are mostly highly cytotoxic CD4+ cells. Importantly. in HIV-1 infected donors under suppressive antiretroviral therapy, we found that these cells are highly enriched for HIV-1 provirus, harbor a clear cytotoxic memory T-cell transcriptome, as well as TCR clonal enrichment. These findings indicate that CD32 remains a promising candidate marker of the HIV-1 reservoir and underline the importance of further investigation of this marker during latent infection.

## Introduction

Fc-gamma receptors (FcγRs) occupy a central role in immunity, mediating important aspects of IgG effector functions. Their prominent expression on macrophages is key to processes like antibody dependent phagocytosis that drives antigen clearance and processing. Historically, among the hematopoietic lineage, T cells were regarded as the exception to the rule as Fc receptors of any kind were not readily detectable on their cell surface (1, 2). Nonetheless, early on, expression of FcγRIIIa (CD16) on CD4+ T cells subsequent to cell activation was reported (3, 4). Reasonably, these studies concluded that the ample evidence for a role of FcγRs on T cells might be due to their strict regulation – fully absent on mature, resting CD4+ T cells, while increased upon activation only (5). In similar lines, the expression of the other class of low affinity FcγRs, FcγRII or CD32, on CD4+ T cells was reported to be upregulated upon cellular activation (3, 6-8), while dim expression on a small percentage of CD4+ T cells was found at baseline (7). Despite that evidence, the role and function of CD32 in CD4+ T-cell biology remains to be defined.

Antiretroviral therapy (ART) effectively suppresses HIV-1 replication and reduces the viral load in plasma to undetectable levels, significantly decreasing HIV-related morbidity and mortality (9). However, ART is not curative and has to be taken lifelong as the therapy fails to eradicate latent HIV-1 reservoirs that reside in various cell types and tissues, ready to reignite active replication if therapy is stopped (10, 11). Persistence of the viral reservoirs in people with HIV-1 (PWH) is the main obstacle to achieving a cure, and the absence of virus-encoded surface markers on the HIV-1 reservoir cells greatly complicates the search for a cure (12). Identification and characterization of host cell markers of HIV-1 reservoirs is thus central to the cure research (13, 14).

Interest to CD32a (FcγRIIa) in the context of HIV-1 cure research skyrocketed due to a major publication identifying this receptor as a long-sought cellular marker of the elusive viral reservoir (15). Several attributes, such as high specificity for quiescent cells combined with an enormous (∼1000-fold) proviral enrichment in CD32+ cells from ART-treated PWH, made this marker so unique and fitting for the concept of the latent reservoir (16). Intensive pursuing research, however, did not reach a consensus, with the only consistent finding across studies being the challenge to purify and define CD32+CD4+ T cells. Two studies characterized trogocytotic T-B cell doublets expressing CD32 (17, 18) and another work suggested that platelet-T-cell aggregates can contribute to the CD32 protein signal (19). In view of these reported tribulations and differences across studies, it is not surprising that also the virological conclusions diverged: while some studies could confirm HIV-1 proviral enrichment in CD32+CD4+ T cells (19, 20), others were unable to demonstrate it (8, 17, 21-23). Our previous report addressed the major pitfalls by developing a two-step MACS-based CD32+CD4+ T-cell purification strategy that drastically reduced the contribution of non-T cells into the CD32+ cell fraction, resulting in a ∼300-fold enrichment for total HIV-1 DNA in this fraction (20). However, that report also did not demonstrate the CD32+ cell purity at the single-cell level. Therefore, a reliable method that would allow the in-depth characterization of CD32 expression on CD4+ T cells is still to be designed.

Against that background, we set out to design a purification strategy that would allow to study CD32+CD4+ T cells in healthy individuals, as well as in PWH. Here we show that a multistep purification procedure enables the isolation of a small population with signs of endogenous, dim CD32 expression. Surprisingly, we could not confirm CD32 upregulation upon T-cell receptor (TCR) stimulation. At a single-cell level, we found cytotoxic memory traits with heightened expression of activation markers. Stimulation confirmed that CD32+CD4+ T cells are a Th1 dominant cytotoxic memory cell subset with TCR clonal enrichment. Finally, we could demonstrate high proviral enrichment, placing CD32 again on the top of the list when compared to other cellular markers of the HIV-1 reservoir.

## Results

### A small subset of CD4+ T cells shows dim CD32 protein expression with traits of endogenous origin

Given the controversy about the nature of CD32 protein expression on CD4+ T cells, we set out to develop a purification strategy, taking shortcomings of previous reports into account. These concern in particular two major and partially intertwined aspects: first, general insufficient purities and second, major contaminations (e.g. T-B-cell doublets, monocytes or platelet-T cell aggregates) in the obtained CD32+ CD4+ T cells (8, 17-19, 22). Undoubtedly, these difficulties are a major contributor to diverging results published on CD32 as a cellular marker of the HIV-1 reservoir. Therefore, a new strategy was needed to generate new insights on the matter. Our procedure consists of three main steps (graphically illustrated in Figure 1A). First, untouched CD4+ T cells are obtained from PBMCs by magnetic-activated cell sorting (MACS), which is followed by two consecutive runs of fluorescent activated cell sorting (FACS). During both FACS sorts, strict exclusion of doublets, monocytes, B-cells, and platelets is applied, as these were hurdles prior attempts had stumbled at (gating strategy is shown in Supplementary Figure 1A). Pursuing that, we observed a median 0.06% of CD4 + T cells dim-positive for CD32 (Figure 1B). As an important side note, usage of Fc blocking reagents did not affect CD32 antibody binding (data not shown), indicating that there is specific antibody reactivity, as reported previously (24).

**Figure 1.**
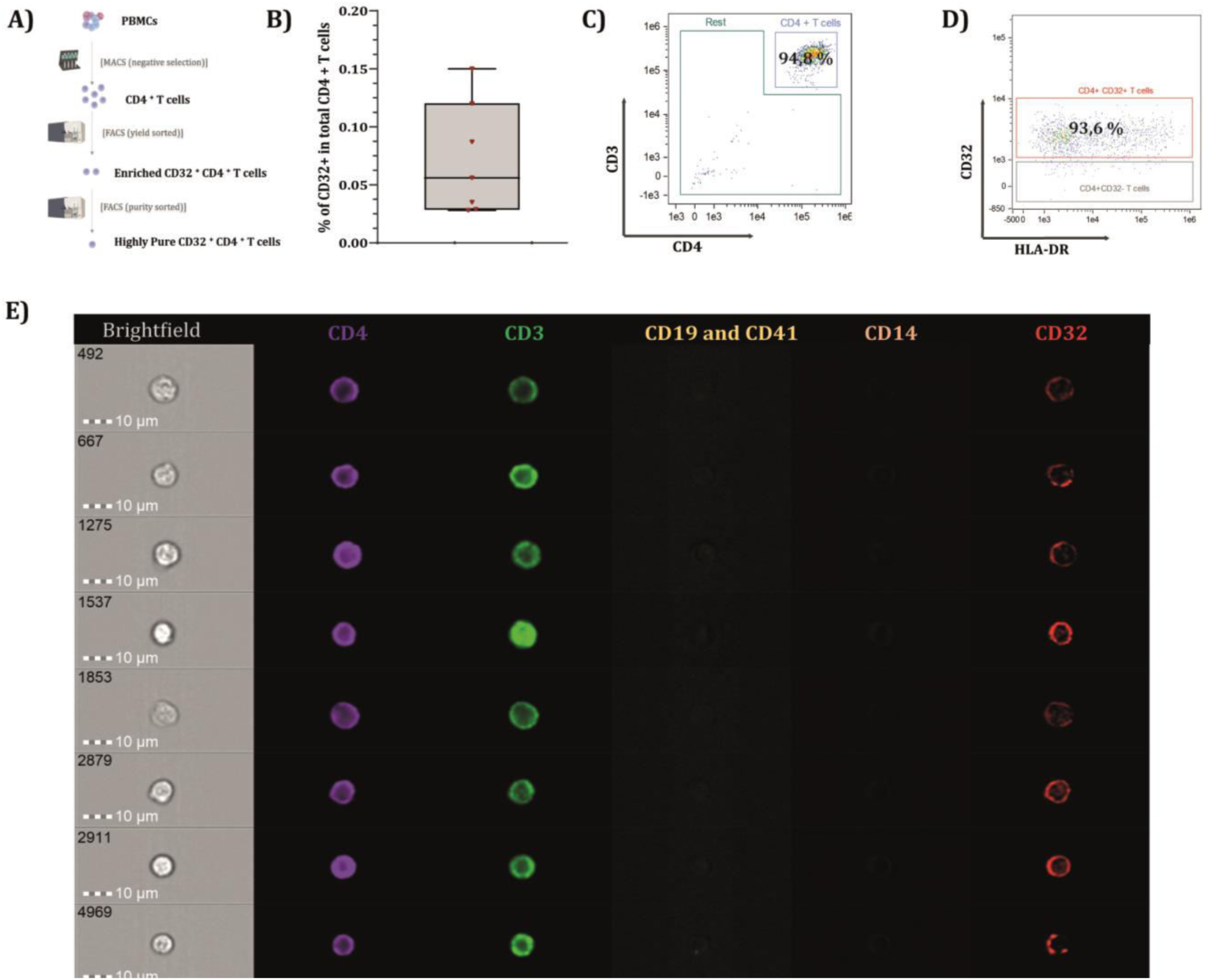
Dim CD32 expression on CD4+ T cells is evinced using a multistep cell sorting strategy. (A) the developed sorting strategy is graphically depicted, consisting of three main steps, MACS isolation of CD4 T cells followed by two consecutive FACS sorts. (B) Median frequencies of bona fide, dim CD32+ CD4+ T cells in periphery are 0.053 % of total CD4 + T cells. (C) The recovered, post-sorted cells show the phenotype of highly pure CD3 + CD4 + T cells. (D) more than 93% of those harbor dim levels of CD32 protein expression. (E) characterization of the staining pattern seen on the highly pure CD32+CD4+ T cells by imaging cytometry suggests endogenous origin due to a homogeneously spread signal over the entire cell membrane.

To validate our new approach and characterize the obtained cell population, post-sorted samples from healthy donors were analysed by imaging cytometry. The analysis showed that more than 94% of all cells were double positive for CD4 and CD3 protein (from here onwards referred to as CD4+ T cells) (Figure 1C). The about 6% of negative cells for CD3 and CD4, were either dying, dead or fragmented cells, while above all, no traces of non-T cell contamination were observed (Supplementary Figure 1B). Importantly, of the obtained CD4+ T cells, more than 93 percent were dim-positive for CD32 (Figure 1D) with no signs of trogocytosis, doublets, or cell aggregates (Figure 1E, channels capturing CD14, CD19 and CD41 expression were consistently negative and bright field shots show single cells). Thus, we report here for the first time a cell purification strategy that enables the pure isolation of CD4+ T cells expressing the CD32 protein from PBMCs. Importantly, the staining pattern (homogeneous signal spread over the cell membrane) suggests the endogenous origin of CD32, as single-cell images showed 90% of CD32^dim^ CD4+ T cells with a uniform stain pattern for the CD32 protein, whereas 10% showed bright punctuate staining, as illustrated in the representative images (Figure 1E). As expected, due to our strict exclusion, the overall proportion of CD4+ T cells expressing CD32 (median, 0.06%) is considerably lower than in most previous studies (6, 8, 21).

### CD32 expression is not profoundly increased upon T-cell stimulation

CD32 exists in two main isoforms: CD32a and CD32b (25), and the anti-CD32 antibody used in this study and most other studies (15, 20) does not discriminate between CD32a and CD32b proteins, because their extracellular domains are very similar (21). In order to determine which isoform of CD32 is expressed on CD4+ T cells, we sought to characterize transcription on ex vivo purified cells from healthy donors by qPCR. On CD32-CD4+ T cells, both mRNAs were detectable, albeit at low levels (Figure 2A). As expected, high levels of the CD32a transcripts were detected on monocyte-T-cell doublets (Figure 2A). Surprisingly, both CD32a and CD32b mRNAs were undetectable on highly pure CD32+ CD4+ T cells (Figure 2A). We speculated that the long processing time of about ten hours could cause loss or degradation of CD32 transcripts, which have been previously shown to be unstable (26). Therefore, we chose next to perform straightforward in vitro activation assays, as several studies had reported significant increases in CD32 mRNA and protein expression under these circumstances (6, 8). This would further allow to answer which isoform(s) of CD32 are expressed by CD4+ T cells. Indeed, MACS-purified CD4+ T cells did increase protein expression about tenfold from 24 hours during activation culture (Figure 2B). CD32a mRNA expression also significantly increased, whereas that of CD32b did not (Figure 2C). Nonetheless, it was surprising to see that the proportion of CD32+ CD4+ T cells after activation was about 10-100 times lower in our hands than in the published literature. We noticed that in prior studies in vitro activation experiments were performed on PBMCs in culture, which we omitted to avoid any potential non-T cell contamination. Consequently, the divergent results are likely rooted in there. Nevertheless, even in our MACS-purified CD4+ T-cell cultures, we found low percentages of non-T cells (Supplementary Figure 2). This made us question, especially using highly sensitive qPCR assays, if the increases observed during activation culture were actually derived from bona fide CD4+ T cells.

**Figure 2.**
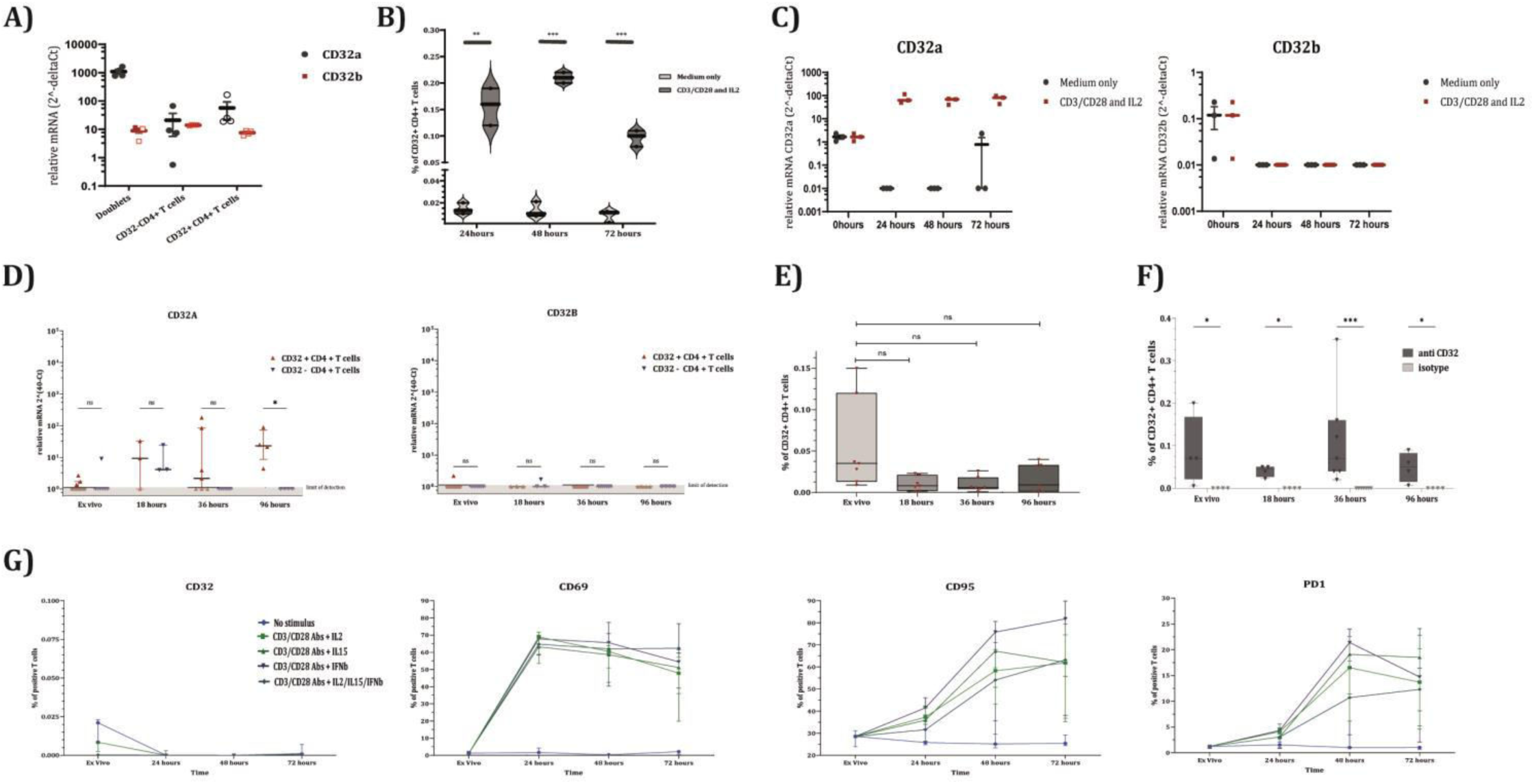
Stimulating highly pure CD4+ T cells does not lead to a marked upregulation of CD32. (A) mRNA of freshly isolated, pure CD32+ CD4+ T cells as well as CD32-CD4+ T cells are undetectable for CD32a (in red) and CD32b transcripts (in dark grey), indicated by non-filled symbols. Control condition, Monocyte-T cell doublets, are as expected clearly detectable for CD32a transcripts. All data is normalized to ribosomal RNA CT values. (B) In culture stimulation of the enriched CD4 + T cells induces significant increases in CD32 protein expression from 24 hours on. (C) Transcripts of CD32a only do significantly increase when CD4+ T cells are stimulated in vitro. (D) Highly pure CD4+ T cells show modest but significant increases in CD32a transcription during activation culture. No increase in CD32b transcription is observed. (E, F) Activation of highly pure CD4+ T cells does not show increases of extracellular as well as intracellular CD32 protein expression. (G) The stimulation of highly pure CD4+ T cells with different stimuli shows clearly that CD32 is not upregulated upon activation whereas early activation (CD69), intermediate activation (CD95) and later activation/exhaustion marker (PD-1) do increase over time. Statistical analysis performed using nonparametric unpaired Wilcoxon rank method.

To address these limitations and shorten the overall processing time (maximizing the chances to detect degradation-sensitive transcripts), we decided to perform a second workflow applying direct on-cell RT-qPCR to omit long sorting. We screened for gene transcripts, as well as extra- and intracellular protein expression levels. To do so, we first purified CD4+ T cells by MACS, followed by a FACS sort leading to >99.5% purity in the final cell culture. Subsequently, triplicates of 100 cells were sorted per donor at every time point according to CD32 expression (CD32+ or CD32-CD4+ T cells) for direct on-cell qPCR measurement. Next to the aforementioned advantages, this workflow also allowed to answer whether CD32 expression is a distinguished feature of a subset of CD4+ T cells, or CD32 is equally expressed by all CD4+ T cells upon stimulation. Interestingly, we could detect very low levels of CD32a isoform only at some time points for both populations. Only at 96 hours we found significant differences between the CD32+ CD4+ T cells and CD32-CD4+ T cells (Figure 2D). However, these minor differences were not paralleled by changes in extra- or intracellular protein expression (Figure 2E and 2F, respectively). CD32b was nearly always undetectable in both populations (Figure 2D). Of note, gene expression levels of the GAPDH reference genes clearly increased over time, indicating the expected activation in culture (Supplementary Figure 2B).

To enlarge our angle, we next assessed if other cytokine stimuli such as IL-15, known to induce a cytotoxic CD4+ T cell profile (27), or the chemokine IFN-β, recently shown to be involved in regulating immune checkpoint expression (28), are involved in CD32 expression, we ultimately tested these cytokines together with CD3/CD28 stimulation. Early, intermediate activation markers, as well as immune checkpoints (CD69, CD95 and PD-1 respectively), did significantly increase over time when compared to unstimulated cells. However, no differences were observed for CD32 (Figure 2G). Taken together, these data indicate that expression of FcγRIIs in CD4+ T cells is not profoundly increased upon cellular activation.

### CD32+ CD4+ T cells have a memory T-cell phenotype with increased expression of activation markers

Given the fact that we succeeded in the isolation of pure CD32+ CD4+ T cells, we next investigated their phenotype. To do so, post-sorted cells from healthy donors were stained with relevant anti memory, activation, immune-checkpoint and chemokine antibodies. The unsupervised analysis of flow cytometry data using the tSNE algorithm was performed on main memory markers (CCR7, CD45RA, CD31, CD95, CD27) and allowed annotation of seven main CD4+ T-cell memory populations (Figure 3A). Next to the general naïve, central memory (CM), effector memory (EM) and terminally differentiated CD45RA re-expressing T cells (TEMRA), we found a memory population (CD45RA-) with high expression levels of CD31 that has not been specifically defined previously. CD31 or Platelet Cell Adhesion Molecule-1 (PECAM-1), is an inhibitory receptor that facilitates TGF-β mediated suppression and is also involved in trans-endothelial migration. Its expression on CD4+ T cells has been associated with naïve, recent thymic migrants (29). We termed this population “transitory memory T cells” as they are of the CD45RA-memory phenotype but show a high level of CD31 expression, allowing them to passage or transit the trans-endothelial junction. Next to that population, we could also clearly distinguish the EM, transitional EM, CM, and terminal CM populations, based on their CD27 expression patterns.

**Figure 3.**
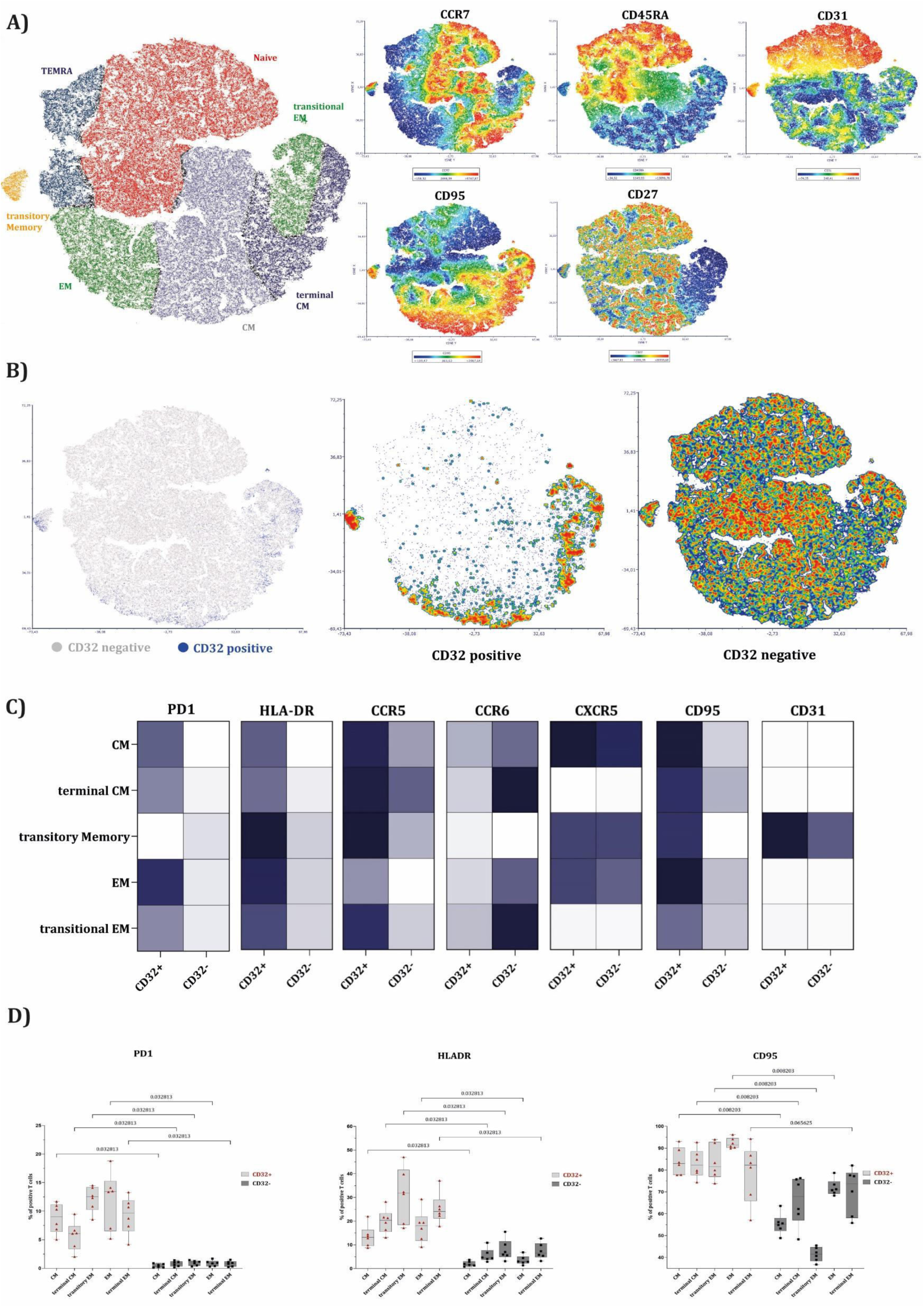
CD32+CD4+ T cells reside in the activated memory compartment. (A) Unsupervised tSNE clustering of flow cytometry data files merged from six healthy donors. Main memory populations are mapped as annotated and colour coded. Calculation of tSNE was based on the expression levels of the CCR7, CD45RA, CD31, CD95, CD27 for which expression levels are shown on the right side. (B) tSNE maps depicting allocation of CD32 + versus CD32– CD4+ T cells. The graph to the left shows distribution of CD32+ CD4+ T cells in blue and CD32– CD4+ T cells in grey. Middle graph depicts density distribution of CD32+ CD4+ T cells only, which mostly densely locate in the memory compartment. Graph to the right shows the density distribution of CD32-CD4+ T cells, which as expected do cluster in all compartments. (C) Heatmaps showing median fluorescence intensities convoluted from six healthy donors per marker of interest for CD32+ versus CD32– CD4+ T cells. Dark blue-grey-white colour legends (from high to low values) are normalized to the expression range of each marker. (D) Percentage of cells expressing PD1, HLADR or CD95 for CD32 + (red symbols on light grey background) versus CD32– (black symbols on dark grey background) CD4+ T cells over different memory compartments. Statistical analysis using Wilcoxon rank test showing p values on the graph.

We were interested to determine where CD32+ CD4+ T cells reside in that memory landscape. Interestingly, >85% of CD32+ CD4+ T cells were of the global memory compartment with barely any TEMRA or naïve cells (Figure 3B and Supplementary Figure 3A). As compared to the CD32-CD4+ T cell population, the relative proportions increased seven-fold for the transitory memory population (1,04 versus 7,56%), three-fold for the terminal EM (5,91 versus 18,24%) and about two and a half-fold for terminal CM (9,28 versus 24,2%) (Supplementary Figure 3). Interestingly, for EM and CM the relative proportions did not change. To adequately compare the expression pattern of activation, chemokine or immune checkpoint markers, we then analysed each of these memory subsets between CD32+ and CD32– CD4+ T cells. The median fluorescence intensity was distinctively higher for CD32+ CD4+ T cells across all memory subsets for all assessed markers, except for CXCR5 and CCR6 (Figure 3C), suggesting increased immune activation and terminal differentiation in CD32+ CD4+ T cells. Significant differences were observed for PD-1, HLADR and CD95 (Figure 3D). This underlines a very distinctive activation profile in CD32+ CD4+ T cells when compared to the respective memory subsets of the global, CD32– CD4+ T cells.

### CD32+ CD4+ T cells have an increased cytotoxic transcriptome and protein signature

Next, we aimed to understand the transcriptional state of CD32+ CD4+ T cells in healthy donors by performing single-cell RNA sequencing. Clustering of the dataset led to the identification of six main populations of CD4+ T cells based on the signature genes they expressed differentially, namely: naïve T cells (SELL, CCR7, LEF1), central memory T cells (CCR7, IL7R, CD28, PASK), T-helper 1 T cells (FOS, HOPX, LYAR, TIMP1, CCL5, IL7R), T-helper 2 T cells (GATA3, KRT1, CAPG, SELL), regulatory T cells (Treg) (FOXP3, IL2RA, SELL) and cytotoxic effector T cells (GZMA, GZMK, NKG7, CST7, PRF1, CD74) (Figures 4A, 4B and Supplementary Figure 4A). Relative frequencies between CD32+ and CD32-CD4+ T cells were 3,1 versus 51,3% for naïve; 16,9 versus 17,1% for central memory; 16,6 versus 14,3% for T-helper 1; 31,2 versus 9,2% for T-helper 2; 27,8 versus 1,2% for Cytotoxic effector and 3,7 versus 6,7% for regulatory T cells, respectively (Figure 4C). The near absence of naïve T cells together with a very high relative increase (about 25 fold), of CD4+ cytotoxic effector T cells identifies a very distinct status of the CD32+ CD4+ T cells.

**Figure 4.**
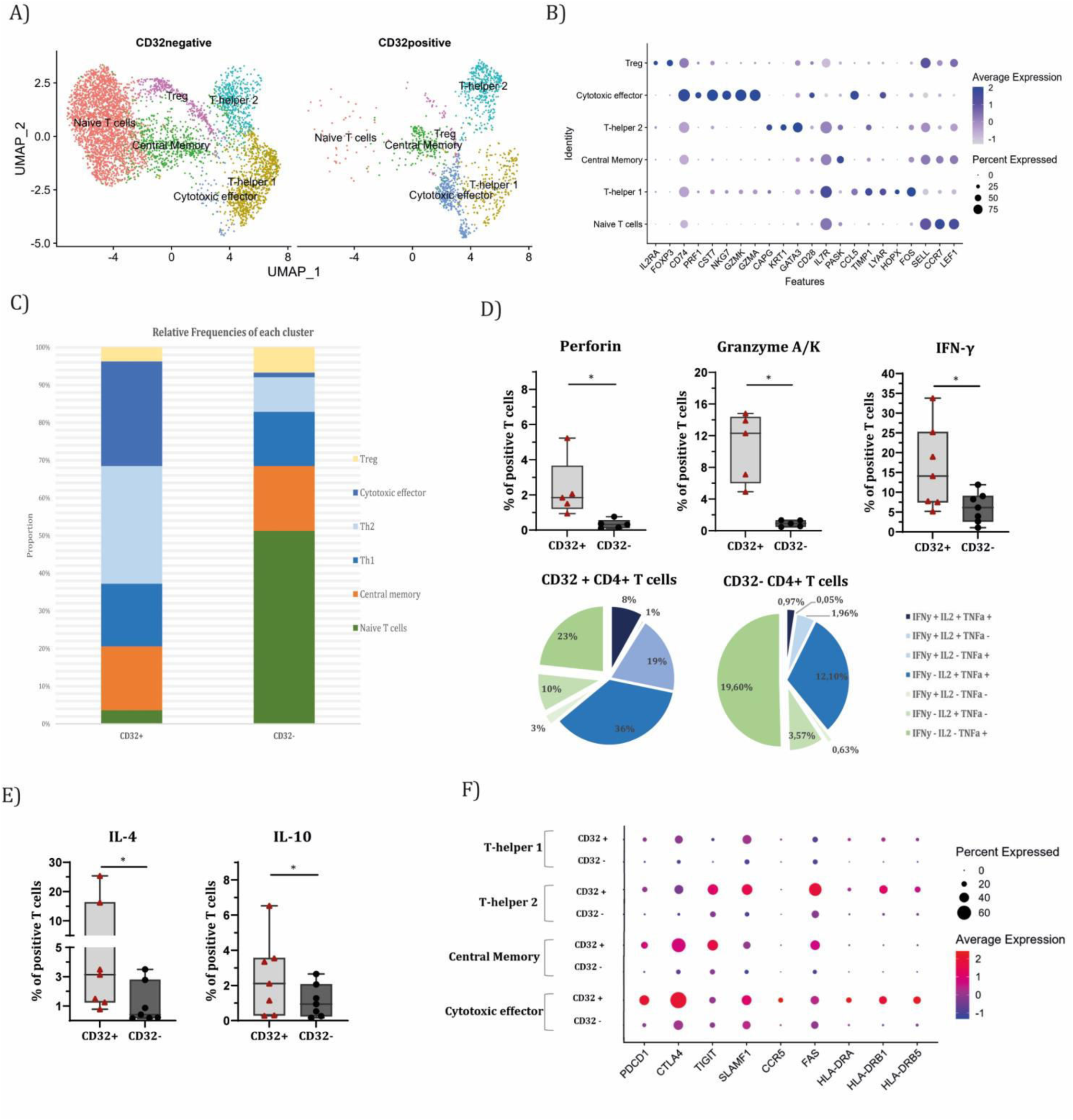
CD32+ CD4+ T cells from healthy donors show a cytotoxic, memory T cell gene expression and protein production profile. (A) scRNA transcriptomics allowed identification of six main CD4+ T cell clusters based on the differentially expressed genes between clusters (shown in B). (C) Relative frequency per cluster for CD32+ and CD32-CD4+ T cells in a stacked bar plot. (D) Intracellular protein staining of Perforin and Granzyme A/K on purified cell populations after 16 hour in vitro reactivation with PMA/Ionomycin as well as cytokines (n=6, non-parametric Wilcoxon rank test with p value cutoff of 0.05). Pie charts show the relative proportion of combinatorial or multifunctional cytokine production profile of Th1-defining cytokines IFN-γ, IL-2 and TNF-α for CD32+ versus CD32-CD4+ T cells. (E) Intracellular protein staining of IL4 and IL10 on purified cell populations after 16 hour in vitro reactivation with PMA/Ionomycin. (n=6, non-parametric Wilcoxon rank test with p value cutoff of 0.05). (F) Gene expression of selected genes over different clusters that either were identified from our prior FACS phenotyping or have an established role in the HIV-1 reservoir field depicted as a dot plot.

We also compared differentially expressed genes across clusters and between CD32+ and CD32-CD4+ T cells. For a general view, we extracted the most significant differentially expressed genes with a minimum log_2_ fold change ≥0,5 for each cluster and plotted them as heatmaps (the naïve and Treg clusters were excluded as those contained <50 cells for CD32+ CD4+ T cells, respectively) (Supplementary Figure 4C). Of the upregulated genes in CD32+ CD4+ T cells (left panels of each heatmap in Supplementary figure 4C), several gene groups can be identified: HLA-I and HLA-II class proteins (HLA-B, HLA-A, CD74, HLA-C), ribosomal proteins (RPS27, RPS29, RPL17, RPL37A), mitochondrial proteins (MT-ND3, MT-ATP6), cell adhesion (ITGB1, ITGB2, CD2, CD99), and cytotoxicity related proteins (GZMA, GZMK). Heightened expression of HLA molecules as well as the mitochondrial genes could indicate cellular stress or even a sign of early stage cell death. However, we could not find significant differences in expression of apoptosis and cell death related genes, albeit a slightly higher but non-significant expression of CASP8 and CFLAR genes in CD32+ CD4+ T cells was observed (Supplementary Figure 4D). The heightened expression of cell adhesion and cytotoxicity related genes suggest a distinct cytotoxic effector gene program in CD32+ CD4+ T cells. Furthermore, four genes coding for ribosomal proteins were significantly upregulated in CD32+ CD4+ T cells, whereas a large group of 24 ribosomal proteins were downregulated. This hints to an overall lower translational activity in CD32+ CD4+ T cells.

For a more stringent look at the differentially expressed genes, we next chose to identify those genes that meet cutoff of log2 fold change ≥1 (Supplementary Figure 4B). We found that for T-helper 1 cell cluster, CD32+ CD4+ T cells expressed higher levels of RPS29 (1.06 fold), SRGN (1.38 fold), ALOX5AP (1.12 fold) and ITGB1 (1.06 fold), and lower levels of IL7R (-1.41 fold). Overall, these genes indicate heightened activation and survival signaling through the IL-7 receptor. For the T-helper 2 cell cluster, we found an increased expression of RPS29 (1.02 fold) and GATA3 (1.19 fold) in CD32+ CD4+ T cells, indicating a more activated and polarized Th2-like transcriptome in these cells. For central memory T cells, SRGN (1.31 fold), PTPRCAP (1.01 fold), ARPC1B (1.01 fold), as well as IL10RA (1.01 fold) were significantly increased, while IL7R (-1.01 fold) was decreased. This gene signature again indicates a more differentiated and activated transcriptome in CD32+ CD4+ T cells, with the exception of IL10RA, which rather indicates a shift into the Th2 or Treg lineage of these central memory cells. Ultimately, in the cytotoxic effector cells, GZMA (1.67 fold), CD74 (1.25fold) and GZMK (1.0 fold) were significantly increased for CD32+ CD4+ T cells, which clearly indicates a heightened polarization into the cytotoxic effector cells. Interestingly, we also found a slightly but non-significantly higher expression of cytotoxicity genes in CD32+ CD4+ T cells of Th1 transcriptome (Supplementary Figure 4E). Of note, we observed higher expression levels of several immune checkpoints (PDCD1, CTLA4, TIGIT), FAS, and HLA-DR classes (Figure 4F), which we for most part already observed on protein level in our FACS phenotyping (Figure 2).

Finally, we assessed if the observed transcriptome profiles are present on the protein level. To do so, we performed short-term in vitro stimulation assays of primary cells. Screening for cytotoxicity associated proteins revealed that CD32+ CD4+ T cells express significantly higher levels of perforin and granzymes A and K (Figure 4D), thus confirming the transcriptome profile. Moreover, CD32+ CD4+ T cells showed significantly increased IFN-γ expression, but also a clear increase in multifunctional profile, defined as co-expression of two or more pro-inflammatory cytokines IFN-γ, IL2, TNF-α by the same cell (64 versus 15,08% for CD32+ and CD32– CD4+ T cells, respectively; Figure 4D). Furthermore, immunosuppressive cytokines that are associated with Th2 functionality were slightly but significantly increased in CD32+ CD4+ T cells, corroborating the transcriptome identified by scRNA-seq (Figure 4E).

### CD32+ CD4+ T cells of PWH are cytotoxic effector T cells with increased expression of CD57, expanded TCR clones, and high enrichment for HIV-1 DNA

Ultimately, we aimed to characterize CD32+ CD4+ T cells in ART-suppressed PWH. Based on single cell transcriptomics, six main clusters were identified (Figure 5A), according to the differentially expressed genes (Figure 5B). Interestingly, next to naïve, central memory and regulatory T cells, we found and annotated three separate clusters, all of which make part of cytotoxic effector T cells. These clusters were distinguished based on differential expression as follows: Th1 effector T cells (GZMK, CXCR3, IFNG-AS1), cytotoxic effector memory (GZMA, NKG7, CST7, PRF1) and cytotoxic terminal natural killer T (NKT) cells (GNLY, FCGR3A, KLRD1, KIR3DL1, GZMH). Overall, the relative frequencies of CD4+ cytotoxic effector T cells (all three populations combined) were <8% for CD32-CD4+ T cells but >80% for CD32+ CD4+ T cells (Figure 5C). Therefore, we observed that the vast majority of CD32+ CD4+ T cells makes part of cytotoxic effector T cells (Figure 5C). Along the same lines, CD32-CD4+ T cells were largely of the naïve compartment (>65%), whereas only a small portion of CD32+ CD4+ T cells (<6%) was of that origin. Looking at the per-cluster quantities, we observed that only the cytotoxic effector memory compartment was present in CD32-CD4+ T cells (cell quantity: 214), while Th1 effector T cells (cell quantity: 6) and cytotoxic terminal NKT cells (cell quantity: 2) were very rare. Conversely, exactly these latter two clusters were highly present in CD32+ CD4+ T cells (173 and 89 cells respectively), underlining the particular, cytotoxic characteristics of these cells. Especially the cluster, which we identified as cytotoxic terminal NKT cells, showed a particular gene expression profile. Next to several NK cell genes, these cells also showed a nearly clonal expression of particular TCR alpha and beta chains (TRAV6-1 and TRBV7-2) (Figure 5B).

**Figure 5.**
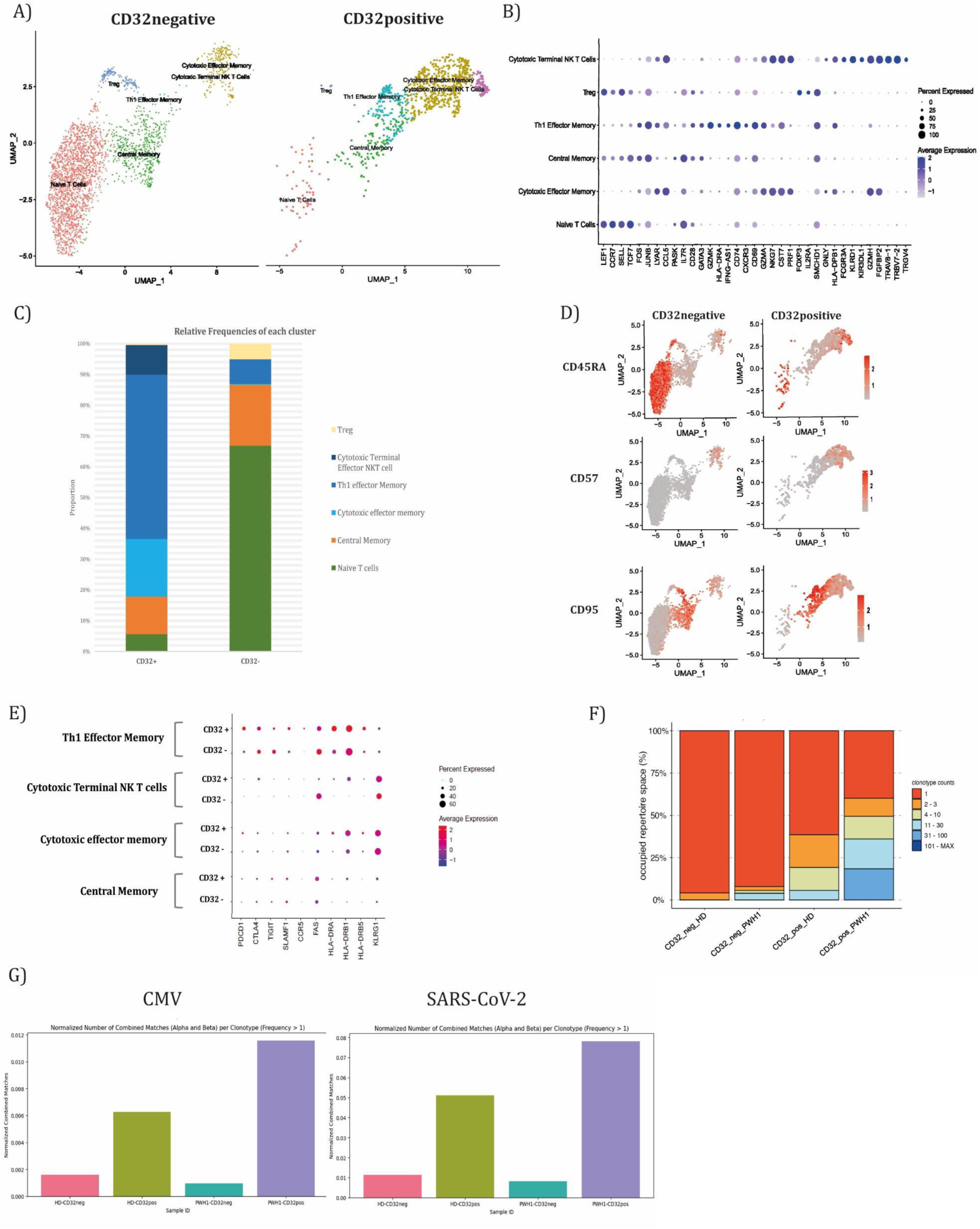
Multiomic analysis of PWH shows clear skewing to cytotoxic memory T cells, clonal TCR enrichment and high enrichment for HIV-1 DNA. (A) scRNA sequenced transcriptomics of HIV-1 infected donor identified six main CD4 + T cell clusters based on the differentially expressed genes between clusters shown in (B). (C) Relative frequencies per cluster for CD32+ and CD32-CD4+ T cells depicted as stacked bar plot. (D) Surface protein expression of tagged antibodies comparing expression levels of CD32-CD4+ T cells with CD32+ CD4+ T cells. (E) Gene expression of selected genes over different clusters that either were identified from our prior FACS phenotyping or have an established role in the HIV-1 reservoir field depicted as a dot plot. (F) Relative frequencies of occupied clonotype repertoire space for CD32+ CD4+ T cells versus CD32-CD4+ T cells in healthy donor (HD) as well as in a person with HIV-1 (PWH). (G) Combined TCR alpha and TCR beta chain matches for clonotypes larger than one shows an increased frequency of CD32+ CD4+ T cells to CMV and EBV epitopes compared to CD32-CD4+ T cells.

Next, we assessed differentially expressed genes of the same clusters but across CD32+ and CD32-CD4+ T cells meeting a cutoff of log2 fold change ≥1. We found a significantly higher expression of some genes (ZFP36, FOS, DUSP1, PPP1R15A, JUN) in CD32+ CD4+ T cells of the central memory cluster, pointing towards a more activated and differentiated state as compared to the general CD32-CD4+ T-cell population (Supplementary Figure 4B). Besides that, we also performed gene set enrichment analysis between CD32+ and CD32-CD4+ T cells of the same cluster. As we had sufficient cell quantities from both (CD32+ and CD32-CD4+ T cells) in the cytotoxic memory compartment only (493 and 214 cells respectively), we included only these results in the study. Globally, the significantly enriched pathways identified cell stress and activation (e.g. HALLMARK HYPOXIA and REACTOME INTERFERON SIGNALING) to be enriched in CD32+ CD4+ T cells, whereas in comparison, gene sets coding for translational activity were enriched in CD32-CD4+ T cells (e.g. TRANSLATIONAL ELONGATION and TRANSLATIONAL INITIATION) (Supplementary Figure 4D). This corroborates the activated, cytotoxic and terminally differentiated status of CD32+ CD4+ T cells, also when compared with CD32-cells from the same cluster.

Next, we performed CITE-seq to assess cell surface protein expression with the antibody-derived tags (ADTs) used during scRNA-seq. Those ADTs were selected based on the gene expression and FACS phenotyping data obtained from the healthy donor data. The selected antibodies were: CD45RA and CCR7 to support the identification of T cell memory status, CD95, an activation marker highly and significantly differentially expressed in CD32+ CD4+ T cells of healthy donors observed in our FACS phenotyping, and CD57, a surface marker known to be expressed by cytotoxic, terminally differentiated CD4+ T cells (30, 31), which, based on our healthy donor transcriptomics, CD32+ CD4+ T cells make part of. As expected, CD45RA surface protein is highly present on naïve T cells. Not surprisingly, we also found increased expression in some of the cytotoxic effector memory T cells as well as the Cytotoxic Terminal Effector NKT cell cluster (Figure 5D), suggesting terminal differentiation of these cells. In similar lines, CD57 was almost exclusively expressed in the cytotoxic effector memory cluster, as well as in the Cytotoxic Terminal Effector NKT cluster, but not in the Th1 effectors (Figure 5D). Interestingly, the cell surface protein expression of CD95 followed a gradient like pattern in CD32+ CD4 T+ cells, from high to low expression (high Th1 effectors, medium in cytotoxic effectors and medium to low in the Cytotoxic Terminal Effector NKT cluster) (Figure 5D), suggesting differences in early/intermediate activation to late stage differentiation. Next, we assessed the expression levels of genes that were linked to HIV-1 reservoirs previously. For these selected genes, we found no differences between CD32+ and CD32-CD4+ T cells over each cluster (Figure 5E).

Next, we performed TCR sequence analysis to investigate how clonotypes distribute over the different cell populations. For this purpose, we combined the TCR data from the healthy and HIV-1 donors into one database. We found clearly expanded clonotypes in CD32+ CD4+ T cells as compared to the CD32-CD4+ T cells (Figure 5F). This difference was more pronounced during HIV-1 infection. Next, in order to probe the antigens those TCR sequences potentially react with, we performed the TCR Match analysis and found that the combined matches for TCR alpha and beta chain were highly increased for CMV and SARS-CoV2 epitopes in CD32+ CD4+ T cells as compared to CD32-CD4+ T cells (Figure 5G). Here again, while the trend was similar, the PWH-derived cells showed pronounced normalized matches. Interestingly, we observed similar but less pronounced trends for EBV and Influenza (Supplementary Figure 5E). Importantly, the TCR match analysis did not show any differences for HIV-1 epitopes (data not shown).

Finally, we purified various CD4+ T-cell cellular subsets based on the expression of CD32 and three other cell surface markers (VLA-4, HLA-DR, PD-1) that have been previously reported to contribute to HIV-1 persistence (32-34). We measured total HIV-1 DNA in subsets of peripheral blood CD4+ T cells from six ART-suppressed PWH that were positive and negative for these markers (Figure 6A). While we observed modest (median, 17.4-fold and 3.9-fold) enrichment for HIV-1 DNA in HLA-DR+ and VLA-4+ CD4+ T cells, respectively, no difference was observed for PD-1 (Figure 6B). For CD32, however, the median and mean enrichment was 151-fold and 284-fold, respectively (Figure 6B), a level that is very close to the enrichment (median, 130-fold; mean, 292-fold) we observed previously in MACS-purified CD32+ CD4+ T cells (20). This finding indicates that pure, cytotoxic, CD32+ CD4+ T cells are highly enriched with HIV-1 provirus.

**Figure 6.**
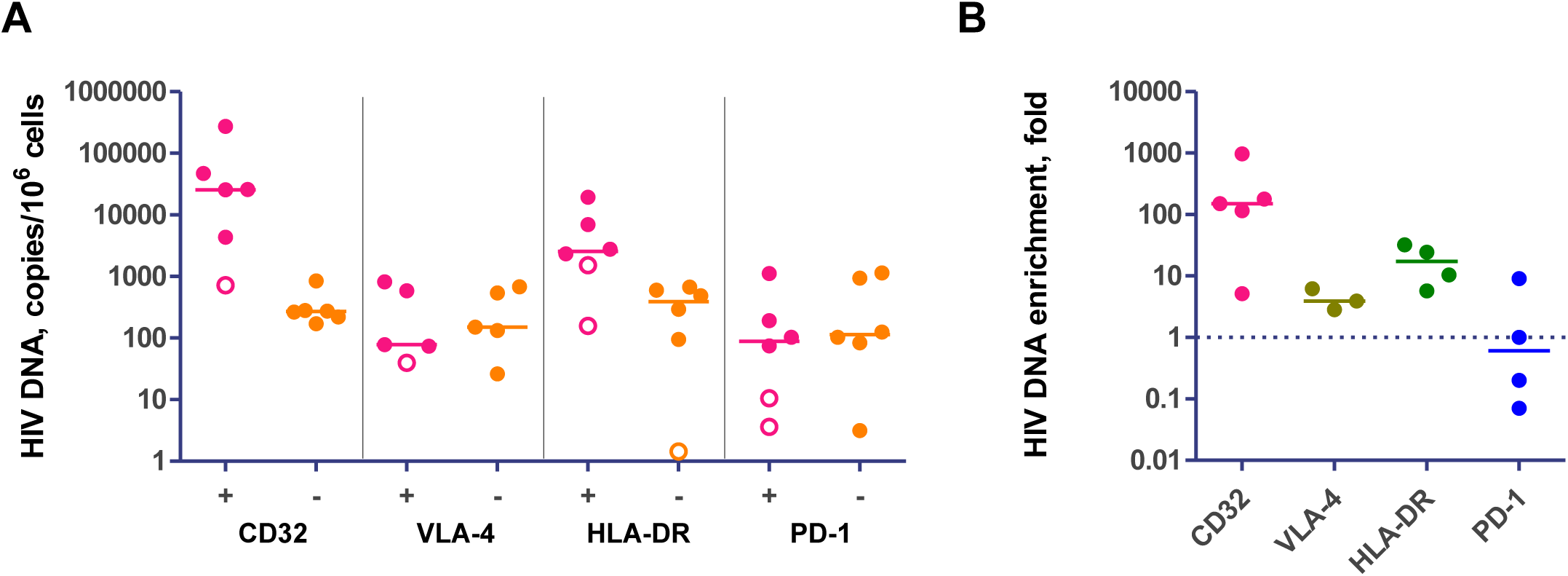
Pure CD32+ CD4+ T cells are highly enriched for HIV-1 DNA. (A) Levels of total HIV-1 DNA in sorted subsets of peripheral blood CD4+ T cells from ART-suppressed PWH (n=6), positive (+) or negative (-) for CD32, VLA-4, HLA-DR, and PD-1. Open circles depict undetectable values, censored to 50% of the assay detection limits. (B) Enrichment values for total HIV-1 DNA on sorted subsets from (A). Enrichment was calculated by dividing the HIV-1 DNA level in the positive subset by its value in the negative subset.

## Discussion

Latent viral reservoirs remain the major hurdle to cure HIV-1 infection. Their scarcity and tissue distribution complicate characterization and monitoring, let alone targeting for eradication (35, 36). Accordingly, research on cellular markers identifying HIV-1 reservoirs is a priority in the HIV-1 cure field (13, 14). It is well established that reservoirs are spread heterogeneously over various CD4+ T-cell subsets and are the source of rebounding virus upon treatment interruption (37, 38). Consequently, it is not surprising that a number of cell surface proteins enrich for HIV-1 provirus, while it appears unlikely that a simple and unique phenotype will allow targeted elimination of all latently infected cells (39-41). Importantly, we now acknowledge that reservoirs reshape in a biased manner during suppressive ART due to the constant immune selection of the reservoir cells and according to the T-cell subsets the reservoirs make part of (42-45). Thus, cellular markers may allow stratification of reservoirs according to the T-cell status and compartment HIV-1 persists in. To this end, CD32 remains a marker of high interest, as the previously reported HIV-1 enrichment in this subset is unparalleled (15, 19, 20). However, to this day we still lack a thorough understanding of CD32+ CD4+ T-cell biology due to tribulations in defining and purifying this CD4+ T-cell subset. To close that gap, we present here an in-depth study of this T-cell population in healthy as well as HIV-1 infected individuals through the establishment of a stringent cell purification strategy.

The nature and origin of CD32 on CD4+ T cells has been subject of a longstanding debate (6, 17-20, 22, 24, 46-48). On the fundamental side, despite some reports, its precise function and role has not been established yet (6, 47). Here we show for the first time that CD32-expressing CD4+ T cells can be purified from PBMCs without signs of non-T-cell-originating contamination or trogocytosis. Importantly, the imaging pattern (homogeneous CD32 signal spread over the membrane) suggests endogenous origin of the CD32 protein. To study its function, we attempted to induce CD32 expression on highly pure CD4+ T cells in vitro using multiple stimuli known to either cause general T-cell activation and expansion (CD3/CD28 beads plus IL2 or PHA plus IL2) or to induce cytotoxic CD4+ T-cell responses (CD3/CD28 plus IL15), as well as antiviral immunity (CD3/CD28 plus IFNb) (27, 28). Surprisingly, none of these stimuli resulted in profound CD32 upregulation on transcriptional or translational levels. This contrasts with published data (6, 8) and hindered a more thorough study of the role of CD32 in CD4+ T-cell biology. We believe that this contradictory finding is caused by the cell types we used. While we observed a clear upregulation in PBMC cultures, less was seen on relatively pure CD4+ T-cell cultures, while ultimately none in highly pure CD4+ T-cell cultures. Intercellular transfer of transmembrane molecules, termed trogocytosis, is therefore the likely cause of the increased CD32 protein levels observed on CD4+ T cells when co-cultured with cell types that highly express CD32, such as monocytes or B cells, in total PBMCs (49-52). Notably, a recent thorough investigation of trogocytotic transfer of CD32 onto CD4+ T cells is in agreement with our observation (53). Consequently, the regulation of bona fide CD32 expression on CD4+ T cells remains an open question. Theoretically, CD32 expression on CD4+ T cells that we observed could have been caused by an earlier trogocytotic transfer, after which protein reshuffling would result in an homogeneous stain pattern. This, however, appears very unlikely, as no other lineage proteins from potential donor cells could be detected (monocytic, B cell, or platelet related proteins: CD14, CD19, and CD41, respectively). Moreover, the dim protein staining pattern, in conjunction with low transcript levels of CD32a in ex vivo isolated CD32+ CD4+ T cells, strongly suggest low endogenous expression, although the stimuli driving this expression still need to be determined.

The phenotype of CD32+ CD4+ T cells has been investigated by several reports before (18, 21, 24, 54), but comes with the major limitation that until now, the isolation of pure CD32+ CD4+ T cells has not been reported. Therefore, we pursued to phenotype these cells after performing the newly developed and validated purification procedure. In general, we found a clear skewing towards the memory T-cell compartment. This trend is in agreement with the fact that proviral persistence is facilitated in memory T cells due to their prolonged survival, proliferation, and maintenance mechanics (55). Furthermore, we observed several additional characteristics. First, we found a high relative increase in the frequency of T cells expressing CD31 in CD32+ CD4+ T cells. Initially, we included CD31 to determine the frequency of naïve, recent thymic emigrant CD4+ T cells (56). However, in CD32+ CD4+ T cells, which are almost exclusively of the memory compartment, a seven-fold higher proportion of cells expressed CD31, which we termed “transitory memory T cells”. While the role of CD31 on memory T cells is fairly neglected, there is actually a substantial amount of data pointing to its diverse functions (57). Of note, CD31 was found to exert protective effects against cytotoxic effector killing, representing an important hint how HIV-1 might persist more effectively in memory cells expressing CD31 (58-60). Next, we could not confirm a prior report (18) that CD32+ CD4+ T cells are of T follicular helper (Tfh) cell phenotype. This divergent observation is probably rooted in the differences in purification strategy, as well as the compartment sampled. Tfh cells, an established site of the HIV-1 reservoir and viral transcription (61, 62), have the natural propensity to form doublets or receive membrane proteins from B cells by trogocytosis, explaining why their detection was linked to CD32 expression in tissues (18). Our strategy did by design strictly exclude any B-cell signal and doublets in order to study bona fide CD32-expressing CD4+ T cells. Furthermore, and in contrast to Thornhill et al. (18), we studied peripheral blood.

Additional phenotypic differences we found were that pure CD32+ CD4+ T cells express higher levels of the immune checkpoint, PD-1, as well as immune activation markers, CD95 and HLA-DR. Noteworthy increases of activation markers have been observed previously in CD32+ CD4+ T cells, however none of these previous studies could demonstrate the purity of the CD32+ CD4+ T cells (8, 21, 24, 54). In HIV-1 latency, the role of PD-1-mediated inhibitory signalling is well documented (63, 64). Furthermore, expression of HLA-DR is appointed to recently divided but chronically activated effector CD4+ T cells. In contrast, CD95, the FAS receptor, appears already during early activation and has a dual function: limiting cellular immune responses by activation-induced cell death, but also exerting costimulatory capacities during primary T-cell activation (65). Interestingly, we indeed found that CD95 surface protein expression is not uniform over CD32+ CD4+ T cells during HIV-1 infection but rather diminishes in late-stage differentiation. Furthermore, a portion of these cells in PWH shows terminal differentiated NKT cell-like characteristics. This finding merits further investigation. In addition, our data shows that using cell surface proteins, CD95 and CD45RA, will allow future works to subdivide the three major clusters in which CD32+ CD4+ T cells reside, thereby allowing an even closer look into where the virus persists. Altogether, our data suggests that exactly this combinatorial profile of (i) activation that fuels cell proliferation, and (ii) expression of immune checkpoints that limits viral gene expression, makes CD32+ CD4+ T cells a highly suitable niche for HIV-1 persistence.

Next to that, a multiomic scRNA sequencing approach allowed us to uncover that CD32+ CD4+ T cells make part of highly cytotoxic effector CD4+ T cells. While cell-mediated cytotoxicity is not a classical role of MHC-II restricted CD4+ T cells, a seminal work found that such cells expand during chronic viral infections as EBV, CMV, and even more so during untreated HIV-1 infection (66). Therefore, expansion of cytotoxic CD4+ T cells is likely linked to a systemic inflammatory condition, as well as antigen-specific responses during a chronic state of viremia (67-69). Remarkably, several single-cell studies over the last years identified cytotoxic CD4+ T cells as important sites of HIV-1 persistence during viremia and suppressive ART, as well as upon latency reversal (70-73). Here we demonstrate that CD32 surface expression identifies a cytotoxic effector population with a high enrichment for HIV-1 DNA. Noteworthy, CD32+ CD4+ T cells show expression of genes and proteins that have been previously identified in HIV-1 reservoir cells, such as PD-1, HLA-DR or KLRG1. The clustering of CD32+ CD4+ T cells in the memory T-cell compartment with cytotoxic gene and protein signatures (increased expression of Granzyme A/K and Perforin compared to CD32-CD4+ cells) observed in this study may provide a clue for the high enrichment for the HIV-1 reservoir in CD32+ cells.

An important question that is still unresolved is how HIV-1 reservoirs are maintained during prolonged suppressive ART. Importantly, antigen-driven clonal expansion of T cells has been found to be a major factor in HIV-1 persistence (74, 75). Through the analysis of TCR sequences, we observed increased clonal expansion in CD32+ CD4+ T cells, which was even clearer during HIV-1 infection. This suggests that these cells respond to antigens that continuously re-challenge immunity and maintain or even expand over time, as reported for CMV (75). Although the current tools inferring antigen specificity from TCR sequences rely on relatively sparse manually curated databases, we identified responses of CD32+ CD4+ T cells to several viral antigens such as CMV, EBV, Flu, SARS-CoV2, and HIV-1. This shows that CD32+ CD4+ T cells have no bias to particular antigens but rather respond to a diverse panel of pathogens. Whether viral clonality and TCR clonality overlap in these reservoirs should be the subject of future research. Finally, it needs to be underlined again that the activated state of cytotoxic CD32+ CD4+ T cells makes them a very particular subset. This status could reflect recently antigenic stimulated cells or even possibly cells that were targeted by residual HIV-1 replication in tissues (14).

In conclusion, we found high HIV-1 DNA enrichment in pure CD32+ CD4+ T cells. While these cells show traits of endogenous CD32 expression, we could not identify the stimulus in vitro. Most interestingly, data from phenotyping and single-cell transcriptomics highlight that CD32+ T cells harbor cytotoxic, activated, and highly differentiated CD4+ T cell characteristics, that are scarce in the general population of CD4+ T cells. Hence, exactly these traits are likely the ideal niche for HIV-1 provirus to persist. Its continuous maintenance during years of suppressive ART might be driven by clonal expansion, as our TCR sequencing data underlines. Future research should focus on detailed longitudinal virological profiling of CD32+ CD4+ T cells reservoirs, which will provide crucial insights and open ways to interfere with their maintenance. Determining the place and role of CD32+ CD4+ cells in the reservoir landscape will be instructive to understand HIV-1 latency.

## STAR Methods

### Donors and sample preparation

All assays (with the exception of total HIV-1 DNA measurements) were performed on freshly isolated peripheral blood mononuclear cells (PBMC). For healthy donors, blood was obtained from standard blood donations supplied as buffy coats by the Sanquin Blood Bank (Amsterdam, Netherlands). HIV-1 infected donors signed informed consent for 100 ml blood sampling drawn at the University Hospital of Liege (Belgium). Characteristics of HIV-1 infected donors are shown in Supplementary Table 1. Samples of HIV-1 infected donors were instantly transported to Amsterdam at room temperature. Blood was immediately processed for downstream analysis. Briefly, buffy coats or full blood was diluted 2 fold in PBS, layered on 15ml Lymphoprep density gradient medium (Stemcell technologies) in 50 ml tubes and then centrifuged at 1200 G for 25 minutes (lowest acceleration and no brake). Next, PBMCs were recovered from the plasma-medium interface and transferred in fresh falcon tubes followed by three washes in PBS at 300 G, 8 minutes to remove platelets. Isolated PBMCs were counted and further processed according to the specific assay.

### T cell sorting from primary PBMCs

Freshly isolated or thawed PBMCs underwent negative magnetic associated cell sorting using the CD4+ T-cell isolation kit (cat. no.: 130-096-533, Miltenyi Biotec,) with LS columns (catalogue no. 130-042-401, Miltenyi Biotec) according to manufacturer’s instructions. Subsequently, CD4+ T cells were counted, spin-pelleted and distributed at a maximum of 25E6 cells in 200ul FACS buffer (PBS supplemented with 0,05% of BSA) per FACS tube (if cell numbers of single donors exceeded that amount, scale-up was done accordingly and stainings were performed in multiple tubes per donor). Next, 20 µl of Human True Stain FcX (Fc receptor blocking solution) (cat. no. 422301, Biolegend, San Diego, California, USA) solution was added and cells were incubated for 10 minutes at 4°C. Then 40 µl of titrated antibody master mix was added per tube, containing anti human CD3 (Clone OKT3, BV510), CD4 (Clone SK3. BV421), CD19 (Clone HIB10, PE), CD14 (Clone M5E2, PE), CD41 (Clone HIP8, PE), CD32 (Clone FUN-2, APC) all Biolegend, LIVE DEAD Near-IR (Invitrogen) and Brilliant Stain Buffer (cat.no: 563794, BD Biosciences). Cells were incubated for 40 minutes at 4°C in the dark. Then samples were thoroughly washed, spin pelleted and resuspended at 15 million cells per ml. Subsequently, cell sorting was performed using a FACS Aria III (BD Biosciences), FACS Fusion S6 (BD Biosciences) or MA900 (Sony Biotechnology) machines. For the first sort, machines were used in yield mode to achieve reasonable pre-enrichment with high cell recovery. Subsequently, to reach the highest possible cell purities, pre-enriched samples were sorted a second time, in purity mode. During all sorting, strict doublet and dead cell exclusion was applied. Furthermore, lineage markers for B cells (CD19), monocytes (CD14) and platelets (CD41) were excluded as dump gates.

### Imaging cytometry

Post sorted cell samples were run through the Amnis® ImageStream®X Mk II imaging flow cytomter (Luminex, Austin, Texas, USA) at 40-fold magnification to characterize the expression pattern of proteins on single-cell level. Data analysis was performed using the IDEAS software version 6.2 (https://www.luminexcorp.com/eu/amnis-imagestream-imaging-flow-cytometer/#software).

### T cell stimulation assays

Post sorted cell samples were cultured in 96 well plates at 1 million cells per ml density in 200 µl of R10 medium. According to the respective assays cells were exposed to different stimuli. For short term stimulation, assessing intracellular cytokine production or cytotoxicity related proteins, cells were exposed to phorbol 12-myristate 13-acetate (PMA) and Ionomycin (both Thermo Fisher Scientific) at 100 ng/ml and 1 µg/ml concentration, respectively, for 18 hours. Transport inhibitors Monensin and Brefeldin A (both Biolegend) were added from 2 hours onwards in culture, according to manufacturer’s instructions (1/1000 final dilution). Multiday CD4+ T-cell stimulations were performed using various compounds at final in culture concentrations as follows: recombinant human IL-2 (Preprotech) - 10ng/ml, recombinant human IL-15 (Biolegend) - 10ng/ml, recombinant IFNbeta (Bio Techne) - 50ng/ml, PHA (Thermo Fisher Scientific) - 5ug/ml. CD3/CD28 dynabeads (Thermo Fisher Scientific) were used according to manufacturer’s instructions. In vitro HIV-1 infection was perfomed by activating purified CD4+ T cells using PHA and IL2 in combination for 72 hours (concentration as given above). Subsequently, cells were harvested, counted, and spinoculated with LAI virus isolate (150 ng p24 virus stock per million CD4+ cells) for 2 hours 1200 G, room temperature, followed by one hour incubation at 37°C. Finally, cells were resuspended and cultured in R10 medium supplemented with IL2. To monitor and verify infection levels, intracellular p24 staining (clone: KC57 in PE) was used.

### Multiplex Quantitative real time PCR for cellular genes

For assessment of cellular gene transcription, two different methods were used. Direct on-cell measurements were performed using the QuantiNova Multiplex one step qRT-PCR reagent kit (Qiagen). Briefly, triplicates of hundred cells per respective subsets were sorted in wells of a 96-well plate containing 9,5 µl of nuclease-free water (Applied Biosystems) per well. Subsequently, cells were lysed through incubation on a heat block for 5 minutes at 74°C and lysates were transferred into qPCR tubes containing the QuantiNova Multiplex reaction mix with one of each Taqman Gene Expression Assays (all Applied Biosystems) for CD3, CD4, CD14, CD19, CD32a, or CD32b (all in FAM), as well as GAPDH (in VIC) as a reference gene. Standard one step RT-qPCR cycling conditions were used on the RotorGene Q thermocycler (Qiagen) thermocycler. The second approach for measuring cellular gene transcription was performed on thousands of cells which were lysed and nucleic acids extracted according to an in-house silica-based method (76). Eluted nucleic acids underwent DNA depletion using the DNA removal kit (Invitrogen) according to the manufacturer’s instructions. Subsequently, reverse transcription was performed using random primers (Qiagen) and SuperScript III reverse transcriptase (Invitrogen). Finally, qPCR was performed with respective Taqman Gene Expression assays (all Applied Biosystems) and Platinum Quantitative PCR Supermix-UDG (Invitrogen) on RotorGene Q thermocycler (Qiagen). Expression values were normalized to ribosomal RNA, measured on the same sample using the TaqMan Ribosomal RNA Control Reagents (Applied Biosystems).

### T cell phenotyping and data analysis

Post sorted samples were spin pelleted and resuspended in 200 µl FACS buffer and 20 µl of Human True Stain FcX solution (Fc receptor blocking solution) (cat. no.: 422301, Biolegend, San Diego, California, USA) was added, followed by 10 minute incubation at 4°C. Then 40 µl of titrated antibody master mix was added, that contained anti human CD45RA (Clone HI100, BUV395), CD69 (Clone FN50, BUV496), CCR7 (Clone 2-L1-A, BUV563), HLADR (Clone L243, BV605), CCR5 (Clone 2D7, BV650), PD1 (Clone EH12.2H7, BV711), CCR6 (Clone G034E3, BV785), CXCR5 (Clone RF8B2, BB515), CD31 (Clone WM59, PE), CD27 (Clone M-T271, PECF594), CD95 (Clone DX2, PECy7), LIVE DEAD Near-IR and Brilliant Stain Buffer. Subsequently, samples were incubated for 40 minutes at 4°C in the dark, then thoroughly washed, spin pelleted and resuspended in FACS buffer followed by acquisition on a BD LSRFortessa™ SORP five-laser flow cytometer. FCS files were cleaned and preprocessed (stream stability, singlets, live cells, non-T cell dump) using FlowJo version 10 and exported for advanced analysis with FCS express software (De Novo Software), applying normalization and transformation pipelines.

### 10x Genomics single cell RNA-seq

Freshly drawn blood samples from healthy donors or ART-suppressed PWH were processed and prepared as described in the section on sample preparation and cell sorting. For PWH, cells were additionally stained with titrated Abs linked to feature barcodes targeting anti human CD45RA (Clone HI100), CCR7 (Clone G043H7), CD95 (Clone DX2), CD57 (Clone QA1704) (all TotalSeqC, Biolegend) and hashtag barcoded Abs (Mixed clones LNH-94, 2M2). After sorting, single cell embedding in gel beads-in-emulsion (GEMs) was performed on a 10x Genomics Chromium Controller, followed by the usage of Chromium Next GEM Single Cell 5’ Reagent Kit v2 (Duals Index) according to manufacturer’s instructions to generate barcoded cDNA. Finally, a separate transcriptome, cell surface protein and V(D)J library were created, followed by sequencing with Illumina NovaSeq S4 sequencer at 150 million reads depths.

### Computational analysis of single cell sequencing data

Raw base call (BCL) files were demultiplexed, aligned, read filtered, barcode and UMI counted performed with the 10x Genomics Cell Ranger analysis pipeline using ‘cellranger count’. Sequence assembly of the V(D)J library was done using ‘cellranger vdj’. Read 1 was assigned 28 base pairs and to identify Illumina library barcodes, cell barcodes and UMIs. Read 2 was mapped to the human reference transcriptome (GRCh38, version 3.0.0) and the CellRanger human V(D)J reference (GRCh38, version 5.0), respectively. Filtering of empty barcodes was performed according to standard CellRanger procedure. Resulting filtered feature-barcode matrices were imported in R (version 4.0.3) using the Seurat package (version 4.0.2). Briefly sequencing files were imported and quality checked (nfeature RNA >800 & nFeature_RNA <2500 & percent.mt <5). Subsequently, normalization, data scaling, principal component analysis and dimensional reduction with uniform manifold projection (UMAP) were performed. Differential gene expression was calculated using the built-in functions. Differentially expressed genes between and within clusters were determined using Seurats built-in functions (FindAllMarkers, FindConservedMarkers, FindMarkers). Gene set enrichment analysis within each cluster between conditions was performed using the fgsea R package (v1.22.0) using the signed p values (i.e. using the average log2 fold changes) obtained using the default the Wilcoxon Rank Sum test from the Seurat FindMarkers function as the rankings. We searched for enrichment in a subset (i.e. HALLMARK, KEGG, REACTOME and BIOCARTA) of the MSigDB (v2023.1.Hs) genesets. TCR sequence analysis was performed using standard workflow of scRepertoire. TCR match was performed by first compiling a database of virus specific TCRs from IEDB (77), VDJb (78) and MCPAS (79) repositories using both MHC-I and MHC-II data. The TCRmatch algorithm (80) (0.97 cut-off) was employed to compare TRA and TRB chain sequences separately against the virus specific TCR library to identify matches. The number of unique matches was corrected for the repertoire size.

### HIV-1 DNA quantification

Nucleic acids were extracted from total PBMCs or sorted CD4+ T cell fractions using the Boom isolation method (16) with the adaptation that two micrograms of Poly-A Carrier RNA (Qiagen) were added per sample prior to the silica binding. HIV-1 DNA was quantified by qPCR using a previously described assay (81). Copy numbers were determined using a 7-point standard curve and normalized to the total cellular DNA (by measurement of β-actin DNA), as previously published (82).

### Quantitation and statistical analysis

Nonparametric analyses (unpaired Mann-Whitney, paired Wilcoxon tests) and correction for multiple comparisons were applied as indicated in figure legends. Statistical analysis was performed using Prism Version 10 (GraphPad Software) or using the built-in functions in respective R packages. All tests were two-sided. P values <0.05 were considered statistically significant.

## Acknowledgements

We acknowledge the participation and commitment of study participants, which made the study possible. We thank Jeroen den Dunnen, Sona Allahverdiyeva, and Lynn Mes for helpful discussions. This study was supported by grant no. 09120011910035 from the Dutch Medical Research Council (ZonMw). AOP acknowledges grant support from amfAR, The Foundation for AIDS Research (grant no. 1110680–77-RPRL), and from Partnership NWO-Dutch AIDS Fonds ‘HIV cure for everyone’ (grant no. KICH2.V4P.AF23.001).

