## Supplementary Material for "Pure CD32+ CD4+ Cells Are Cytotoxic Memory CD4+ T Lymphocytes Highly Enriched for HIV-1 DNA"

A)

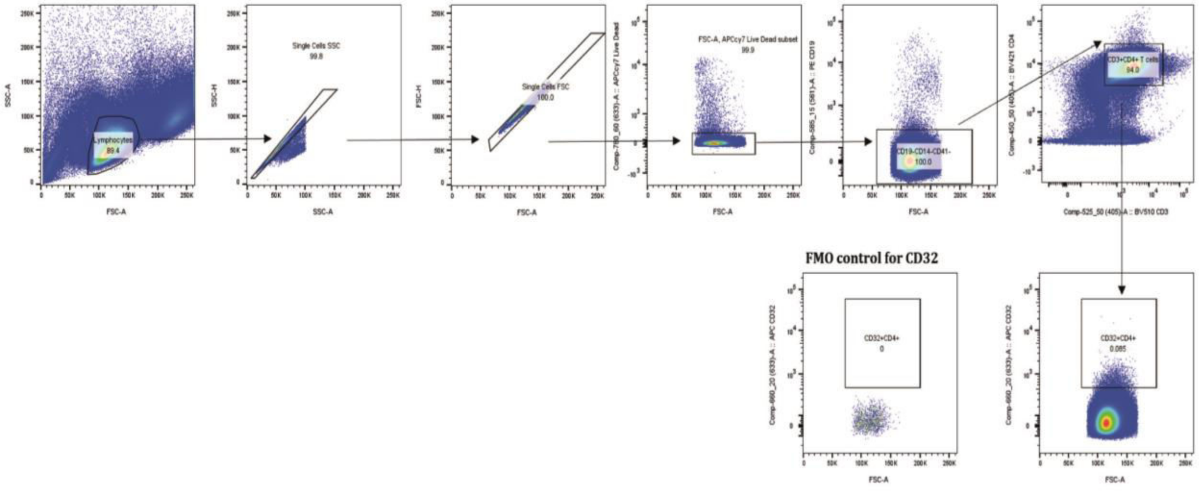


B)

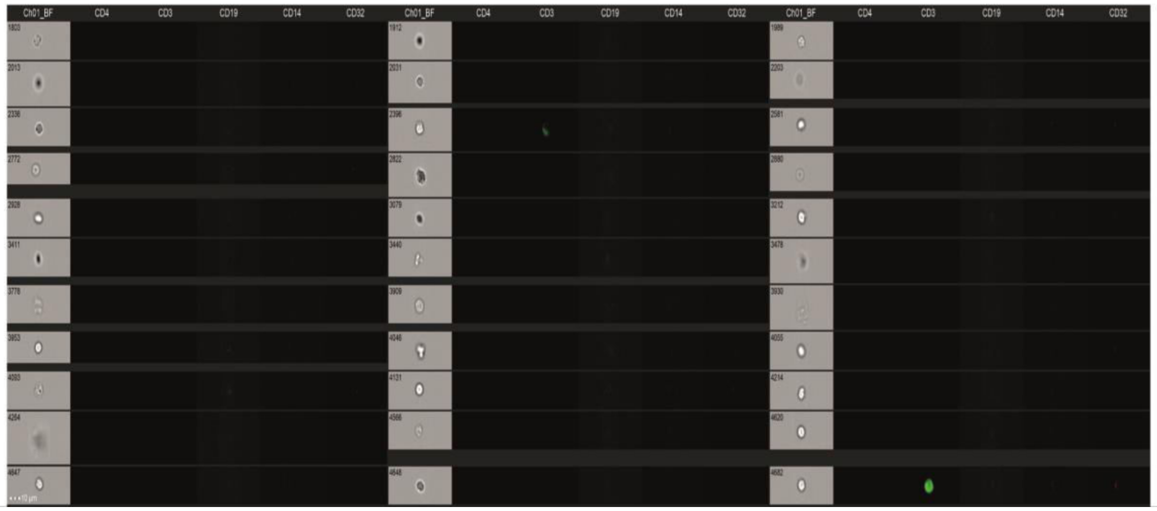


Supplementary figure 1

A)

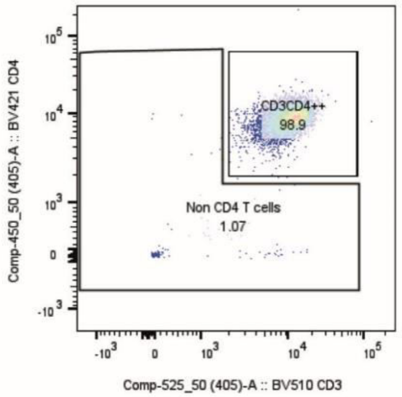


B)

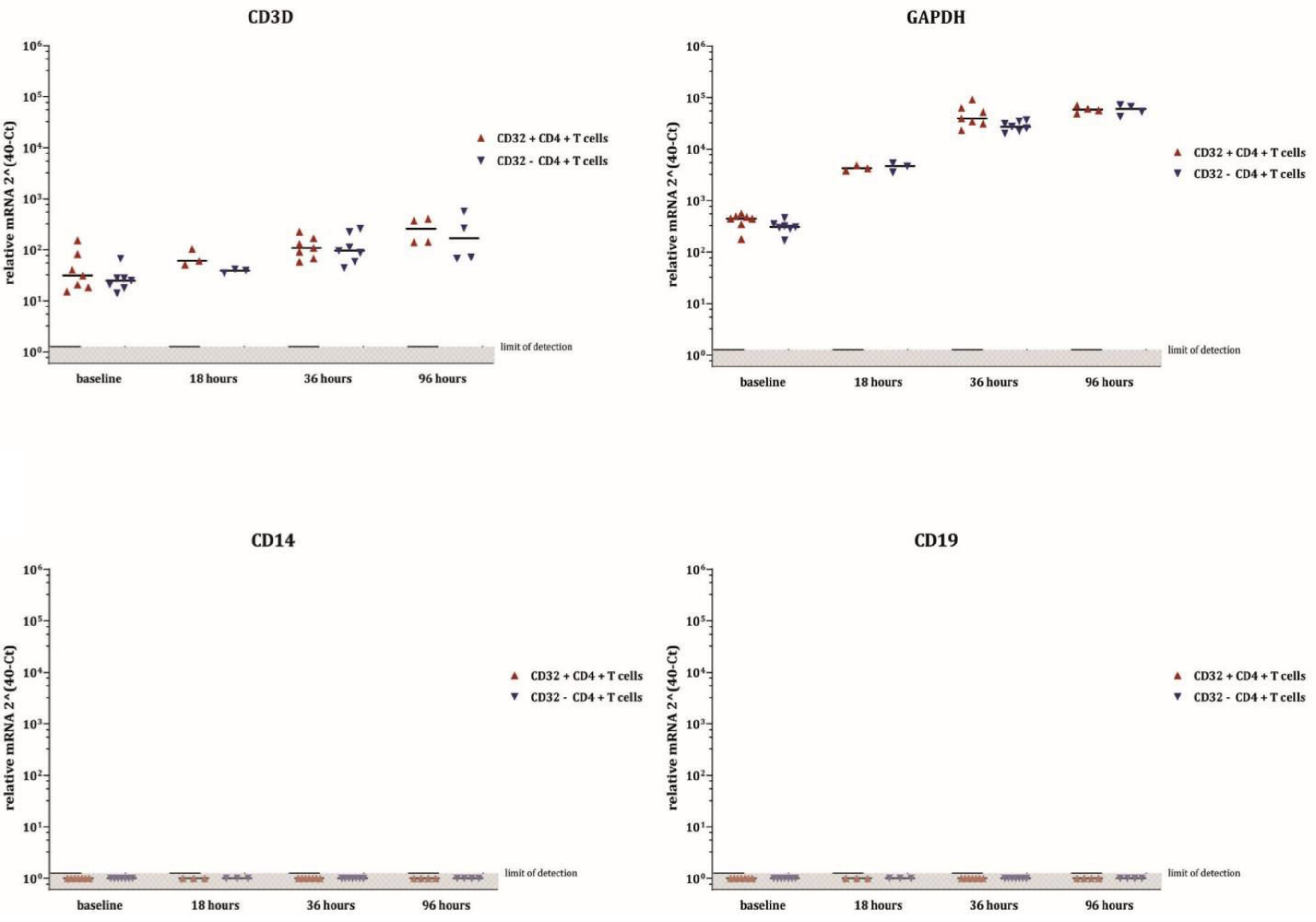


Supplementary figure 2

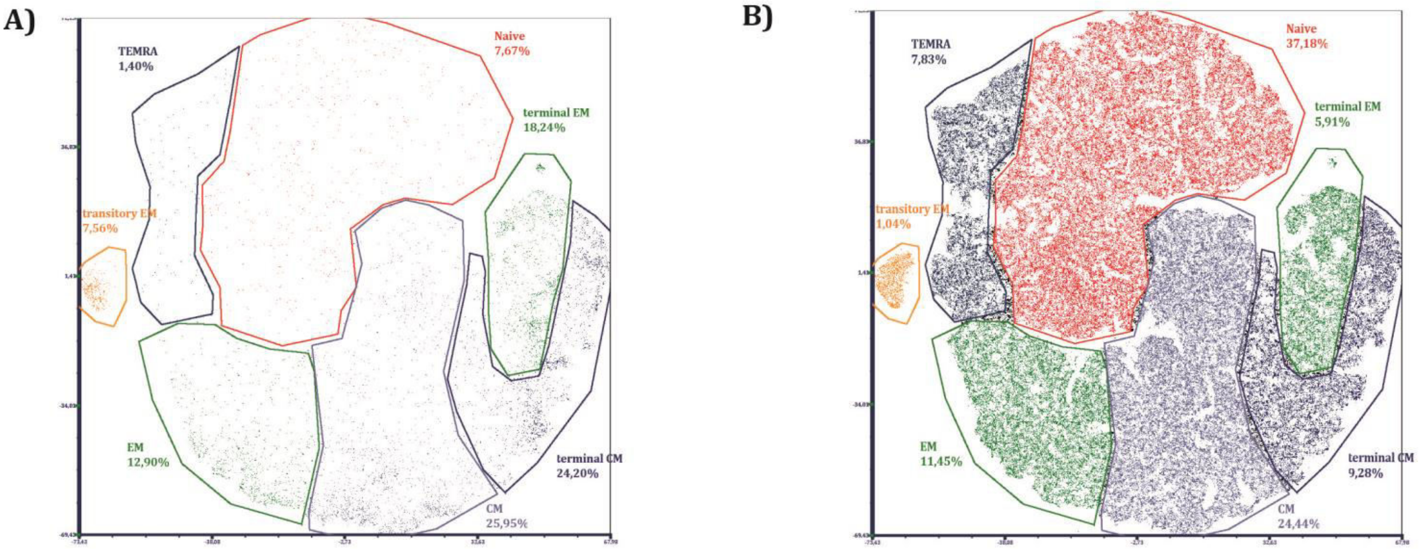


Supplementary figure 3

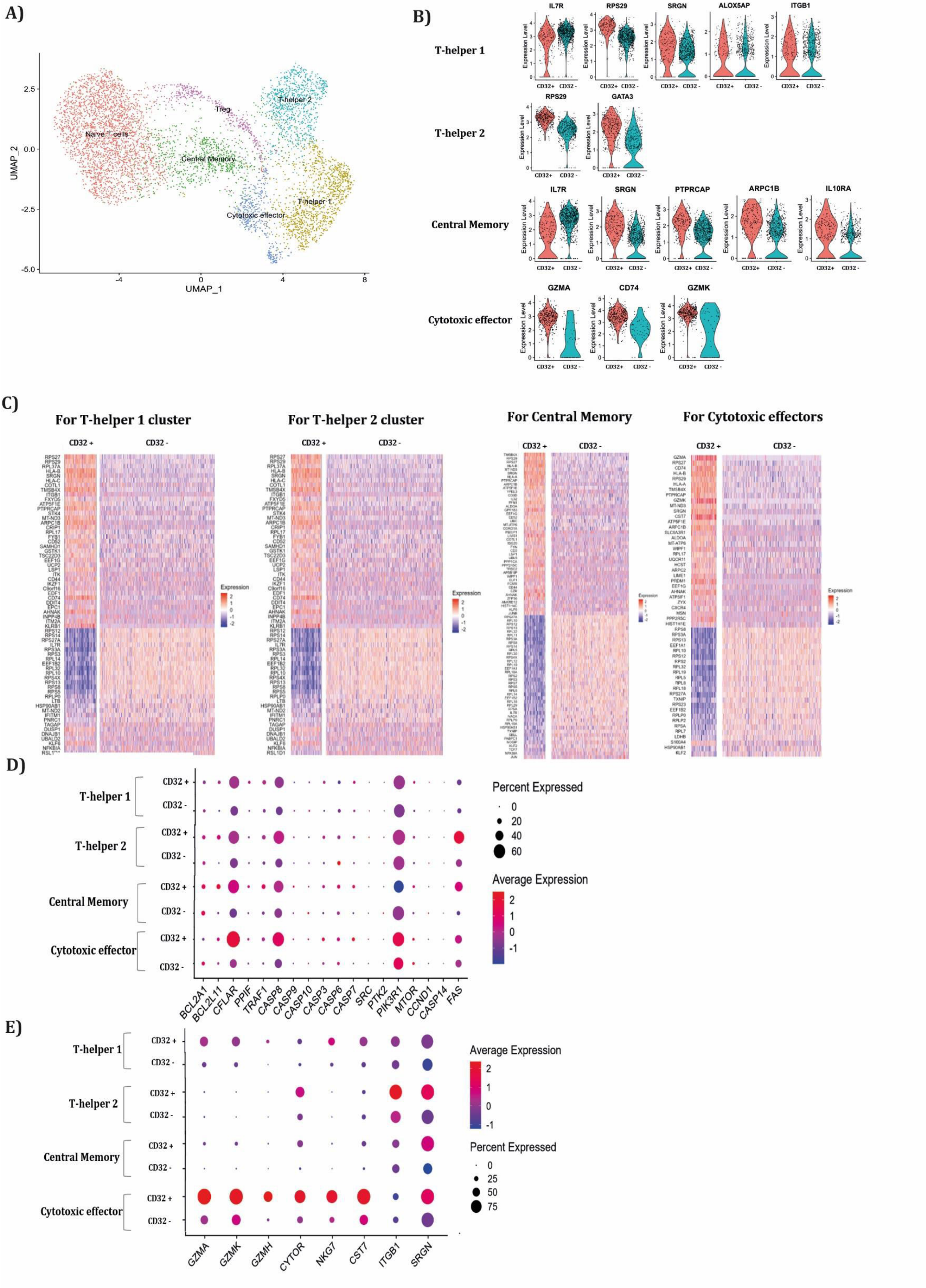


Supplementary figure 4

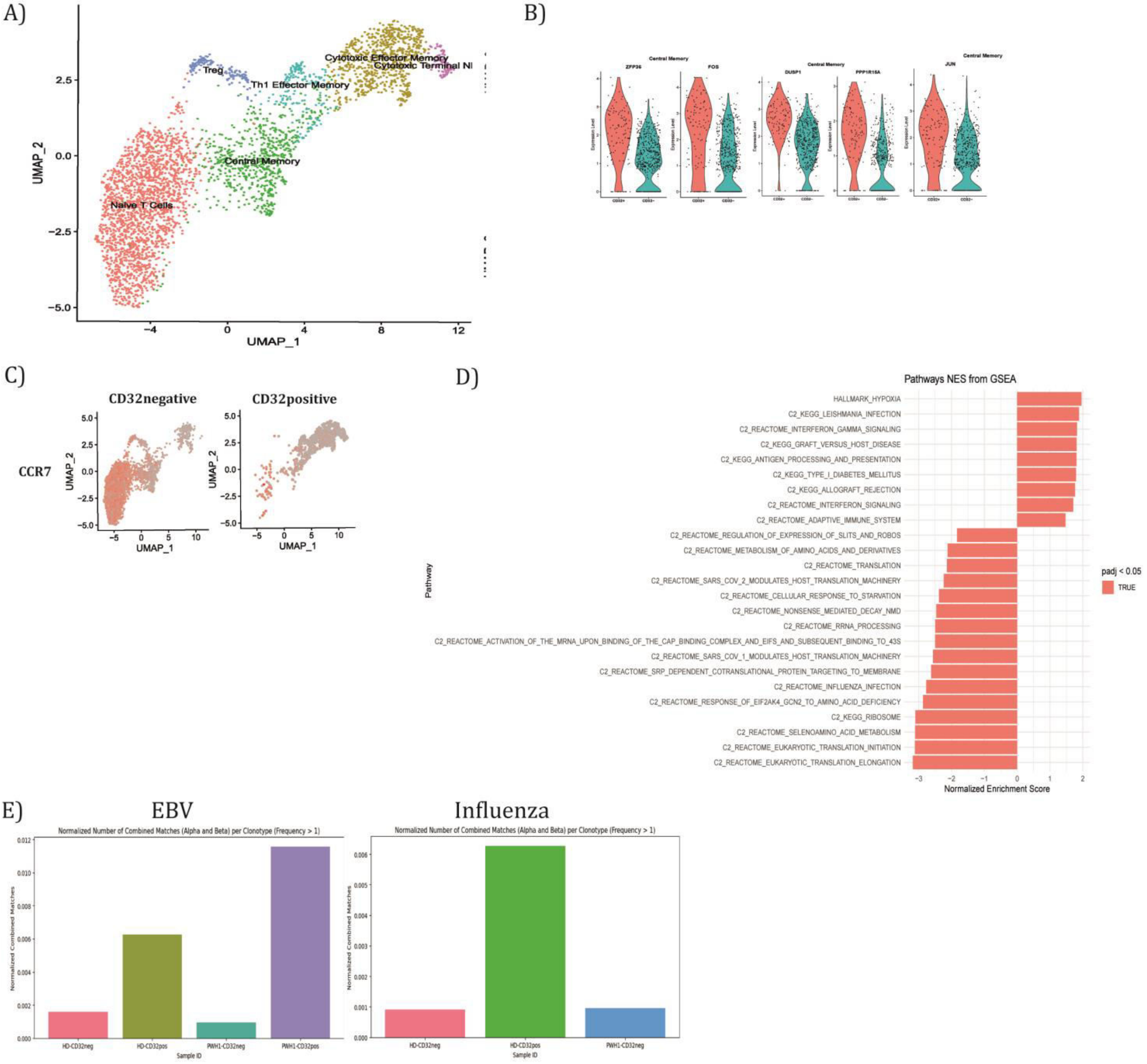


Supplementary figure 5

**Supplementary figure legends**

**Supplementary figure 1. Cell sorting gating strategy and ImageStream gallery of remainder non-CD4+ T cells after double sorting.** (A) Gating strategy applied during cell sorting after MACS isolation of CD4+ T cells. Gates are hierarchically concatenated starting (lymphocytes, SSC and FSC doublet exclusion, live cells, triple negative for CD19, CD14, and CD41 (to prevent B cell, monocyte or platelet contamination, respectively), CD3+ CD4+ T cells and then on CD32 (FMO gate shown to ensure background control). (B) Image gallery displaying non-CD4+ T-cell events in the post double sorted samples of CD32+ CD4+ T cells to investigate for potential contaminating cell populations.

**Supplementary figure 2. Remaining non-CD4+ T cells and gene expression during culture.** (A) Presence of non CD4 T cells in double MACS purified PBMCs shows ≥ 1% of contamination. (B) Gene expression using direct on cell RT-qPCR of cell lineage transcripts CD3D, CD14, and CD19. The reference gene GAPDH was assessed to monitor metabolic activity during stimulation culture. Clear increases over time corroborate TCR-mediated activation.

**Supplementary figure 3. Relative abundance of CD32+ versus CD32- CD4+ T cells across memory populations in tSNE-processed FACS phenotyping data.** (A) CD32+ CD4+ T cells reside with more than 85% in the memory T cell compartment. (B) The CD32- CD4+ T cells show a distribution as expected with nearly 40% of naïve and about 60% of memory T cells.

**Supplementary figure 4. Single-cell RNA sequencing profile of CD32+ versus CD32- CD4+ T cells in healthy donor.** (A) Clustering of total CD4+ T cells using UMAP identifies six main clusters annotated based on the differential expression of canonical subset genes. (B) Significantly differentially expressed genes across clusters between CD32+ and CD32- CD4+ T cells with a cutoff of 1 average $log_2$ fold change depicted as violin plots (Wilcoxon Rank sum test, all adjusted p values below $10^{-8}$). (C) Significantly differentially expressed genes between CD32+ and CD32- CD4 + T cells over the four relevant clusters (cutoffs, minimum log2 fold change of 0.5 and expression in at least 50% of cells in both conditions). (D) Dot plot of genes involved in cell death pathways over cell clusters. (E) Expression of cytotoxicity-related genes over cell clusters.

**Supplementary figure 5. Single-cell RNA sequencing profile of CD32+ versus CD32- CD4+ T cells in HIV-infected ART-suppressed donor**. (A) Clustering of total CD4 + T cells using UMAP identifies six main clusters annotated based on the differential expression of canonical subset genes. (B) Significantly differentially expressed genes across clusters between CD32+ and CD32- CD4+ T cells with a cutoff of 1 average $\log_2$ fold change depicted as violin plots (this cutoff resulted in the fact that only the clusters of central memory T cells was found) (Wilcoxon Rank sum test, all adjusted p values below $10^{-8}$). (C) ADT feature plot for CCR7. (D) Gene set enrichment analysis for the cluster of cytotoxic memory T cells reveals nine gene sets enriched in CD32+ CD4+ T cells while sixteen unenriched as compared to CD32- CD4+ T cells. (E) TCR match results for alpha and beta chain matches to EBV and Influenza databases showing normalized repertoire space per clonotypes >1.

**Supplementary table 1.** Characteristics of HIV-1 infected donors.

| Donor | Sex at birth | Age, years | Plasma viral load, copies/ml | Years since first positive HIV-1 serology | ART regimen | CD4 count, cells/mm$^3$ |
|---|---|---|---|---|---|---|
| D1 | M | 39 | 37[a] | 7.80 | DTG/3TC | 1580 |
| D2 | M | 66 | <20 | 13.40 | BIC/FTC/TAF | 1160 |
| D3 | F | 40 | <20 | 8.23 | DTG/3TC | 640 |
| D4 | M | 65 | 70[a] | 14.21 | CAB/RPV | 1080 |
| D5 | M | 43 | 43[a] | 6.67 | EVGc/FTC/TDF | 1340 |
| D6 | M | 53 | <20 | 11.99 | DTG/3TC | 770 |
| D7 | M | 44 | <20 | 13.34 | DTG/3TC | 1240 |
| D8 | M | 67 | <20 | 27.36 | DTG/3TC | 870 |

[a] Represents an isolated "blip"; plasma viral load measurements before and after were <20 copies/ml.
